# Dilatable DNA origami nanopores as nuclear pore mimics

**DOI:** 10.64898/2026.09.03.749129

**Authors:** Eva Bertosin, Bert van Herck, Anders Barth, Shuo Wang, Cees Dekker

**Author notes:** Equally contributing (Eva Bertosin, Bert van Herck).

## Abstract

The nuclear pore complex (NPC) is a massive protein system that controls all nucleocytoplasmic transport via a dynamic network of intrinsically disordered proteins called FG-Nups. We developed an NPC-mimicking DNA origami nanostructure that self-assembles into a tetrameric, octagonal nanopore with an inner diameter that, through addition of DNA oligonucleotides, can be tuned in a user-defined fashion from 57 to 66 nm – mimicking the contracted and dilated states of NPCs. We demonstrated that these origami nanopores can dilate within minutes and recontract at slower timescales. Both contracted and dilated pores can be inserted into lipid bilayers, where they maintain their conformation and allow transmembrane transport of fluorescent proteins. With outer diameters up to 87 nm, these are, to our knowledge, the largest DNA origami nanopores inserted into lipid membranes. Functionalization of the pores with the FG-Nup Nsp1 reduces non-specific cargo diffusion across the lipid membrane while facilitating translocation of the transport receptor Kap95, demonstrating transport selectivity much like biological NPCs. The work establishes a size-tunable platform to dissect nuclear transport mechanisms, with potential applications for engineering programmable artificial channels.

## Introduction

The nuclear pore complex (NPC) [1, 2] is a massive protein system that spans the nuclear double membrane of eukaryotic cells [3–5] to control all molecular transport into and out of the nucleus. With a mass of ∼110 MDa in humans [6], it assembles from ∼35 different proteins present in multiple copies, with a total of almost 1000 protein subunits [2]. The NPC comprises an octagonal, ring-like structure of nucleoporin proteins (Nups) that is filled with intrinsically disordered proteins (FG-Nups) rich in phenylalanine-glycine repeats [7]. These FG-repeats establish a selective permeable barrier between the nucleoplasm and the cytoplasm, allowing only small molecules with molecular weights below ∼40 kDa to diffuse across the NPC [8, 9]. In contrast, larger macromolecules can cross the nuclear barrier only when bound to nuclear transport receptor (NTR) proteins such as the karyopherin (Kap) family [10], which recognize nuclear localization sequences (NLS) in cargos [11]. Remarkably, upon interacting with the FG-repeats of FG-Nups, large cargo-loaded NTRs effectively cross the mesh on a millisecond time scale [12–16]. Recent cryo-electron microscopy (cryo-EM) studies showed that NPCs are not static structures but they can dilate or contract in size depending on the state of the cell [17, 18], with an inner lumen diameter that can vary in response to variations in membrane tension [2], from 49 nm under stress to 69 nm under hypo-osmotic conditions (Fig. 1) [17, 18].

**Figure 1:**
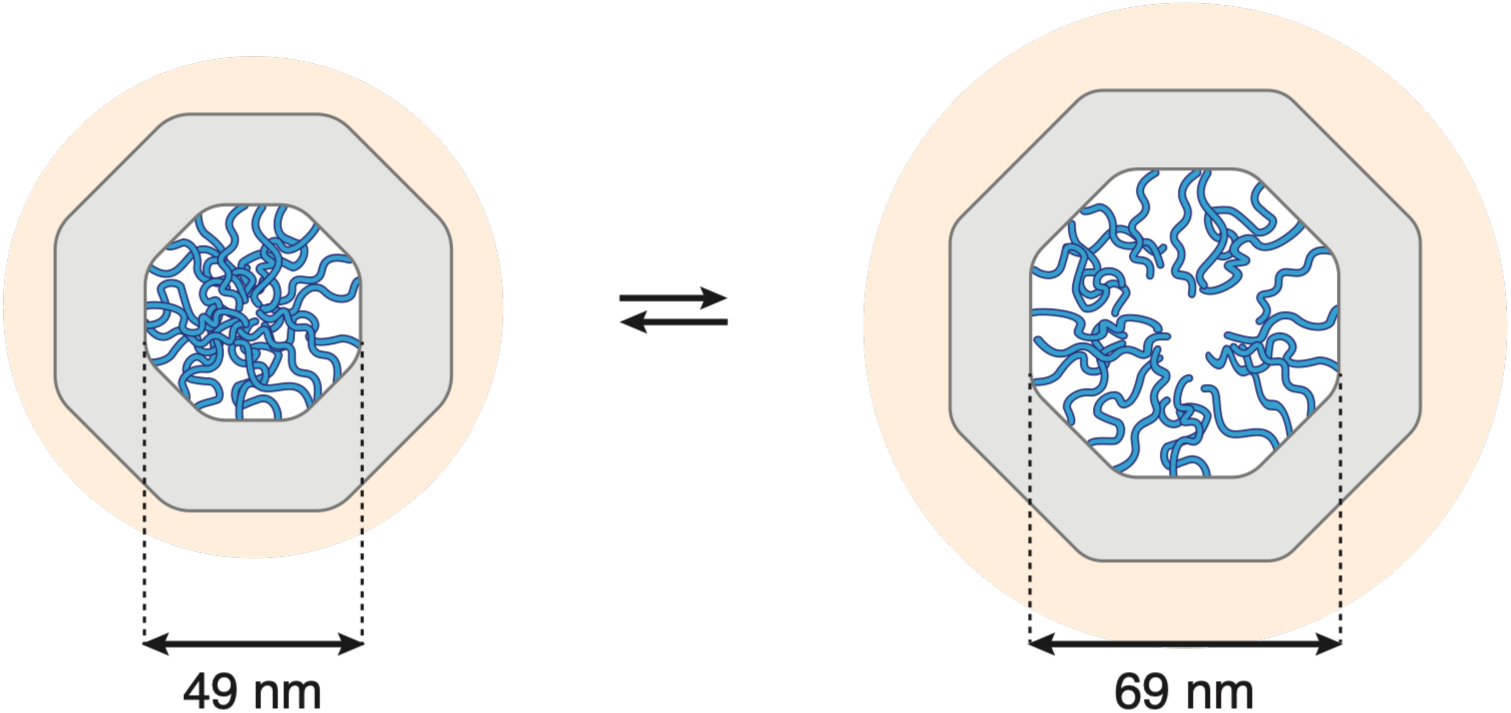
The Nuclear Pore Complex. Schematic representation of the contracted (left) and dilated (right) biological NPC. Scaffold Nups are represented as a grey octagon, FG-Nups as blue lines, and nuclear envelope as the peach-colored area. Dimensions from [17].

Due to the large number, intrinsic disorder, and fast dynamics of the NPC components, it has remained very challenging to dissect the biophysical mechanism underlying nuclear transport *in vivo* [19–22]. Moreover, the effects of the NPC dilation and contraction have hardly been explored. To overcome the challenges of examining NPCs *in vivo*, several bottom-up *in vitro* approaches have been developed [23, 24]. Early examples comprise FG-Nup-coated SiN solid-state nanopores through which the translocation of NTRs could be studied using ion current measurements [25, 26], while optical monitoring with zero-mode waveguides was also explored [27–30]. An alternative to nanopores in inorganic materials is to use organic-based constructs. Indeed, DNA nanotechnology has emerged as one of the most promising methods to build biomimetic NPCs from the bottom up. Specifically, ‘DNA origami’, in which structures are built from a single-stranded (ss) circular scaffold that is stabilized in the desired shape by means of hundreds of shorter oligonucleotides [31–33], is a powerful method that allows the construction of 2D or 3D nanostructures with custom shapes and sizes [32–34]. DNA origami nanostructures have been used to mimic various aspects of NPCs, e.g., the presence of different types of FG-Nups or the ability to insert into lipid bilayers [35–42]. However, so far, these nanostructures were static and did not recapitulate the ability of the NPC to dilate and contract.

Here, we built a DNA origami nanostructure that can dilate and contract similarly to the NPC. These origami structures self-assemble into large tetrameric nanopores with octagonal symmetry, with inner diameters spanning from 57 to 66 nm (outer diameters from 74 to 87 nm, Table S1). We demonstrated that these ultrawide octagons can be inserted into lipid bilayers, making these, to our knowledge, the largest DNA origami nanopores inserted into membranes. As shown via fluorescence lifetime imaging, the nanopores maintain their contracted or dilated state when embedded into membranes. Using TEM and single-molecule FRET experiments, we proved that the nanopores can dilate and contract in a controlled way upon administering DNA oligonucleotides (oligos). We probed the transmembrane molecular transport properties of the nanopores via influx of fluorescent proteins, and demonstrated that it depends on the octagon diameter. To mimic the selective transport abilities of the NPC, we decorated the inner lumen of the nanostructures with up to 120 FG-Nups and inserted the functionalized structures into lipid bilayers of a giant unilamellar vesicle (GUV), thus constituting an artificial nucleus, to study transmembrane protein transport. Functionalizing nanopores with Nsp1 conferred selective permeability as the diffusion of an inert protein into the GUV lumen is restricted while Kap95 underwent facilitated transport. The work marks a step towards building controllable mimics of the NPC that can recapitulate its selective transport properties. Future studies will use this platform to investigate the biophysical principles of protein transport through the NPC, with potential applications in the field of synthetic biology.

## Results

### Building tetrameric octagonal origami rings

Inspired by the NPC, we designed a nanopore with octagonal symmetry that is aimed to be embedded within a lipid bilayer membrane. We note that it is practically not feasible to replicate the very large dimensions of biological NPCs with a single DNA origami structure, as the scaffold length is limited to ∼8000 nucleotides (nt), making the final structure inherently thin and floppy and leaving an insufficient number of anchor points for further functionalization. Hence, we designed a monomeric origami with a shape corresponding to one-quarter of a regular octagon (Fig. 2A). In top view, the structure comprises a long inner edge (∼24.5 nm) flanked by two shorter edges (∼12 nm each). The angle between the long edge and the shorter ones is 135°, consistent with the geometry of a regular octagon (Fig. 2A, right). In a cross-sectional view, the origami comprises a transmembrane vertical part connected to a wider horizontal flap element (Fig. S1). The transmembrane part forms the membrane-spanning region, while the flap extends laterally above the membrane to guide the insertion of the structure into lipid bilayers [43]. Additionally, the monomer interfaces are designed such that one surface is shape-complementary to the opposite interface (Fig. S2).

**Figure 2:**
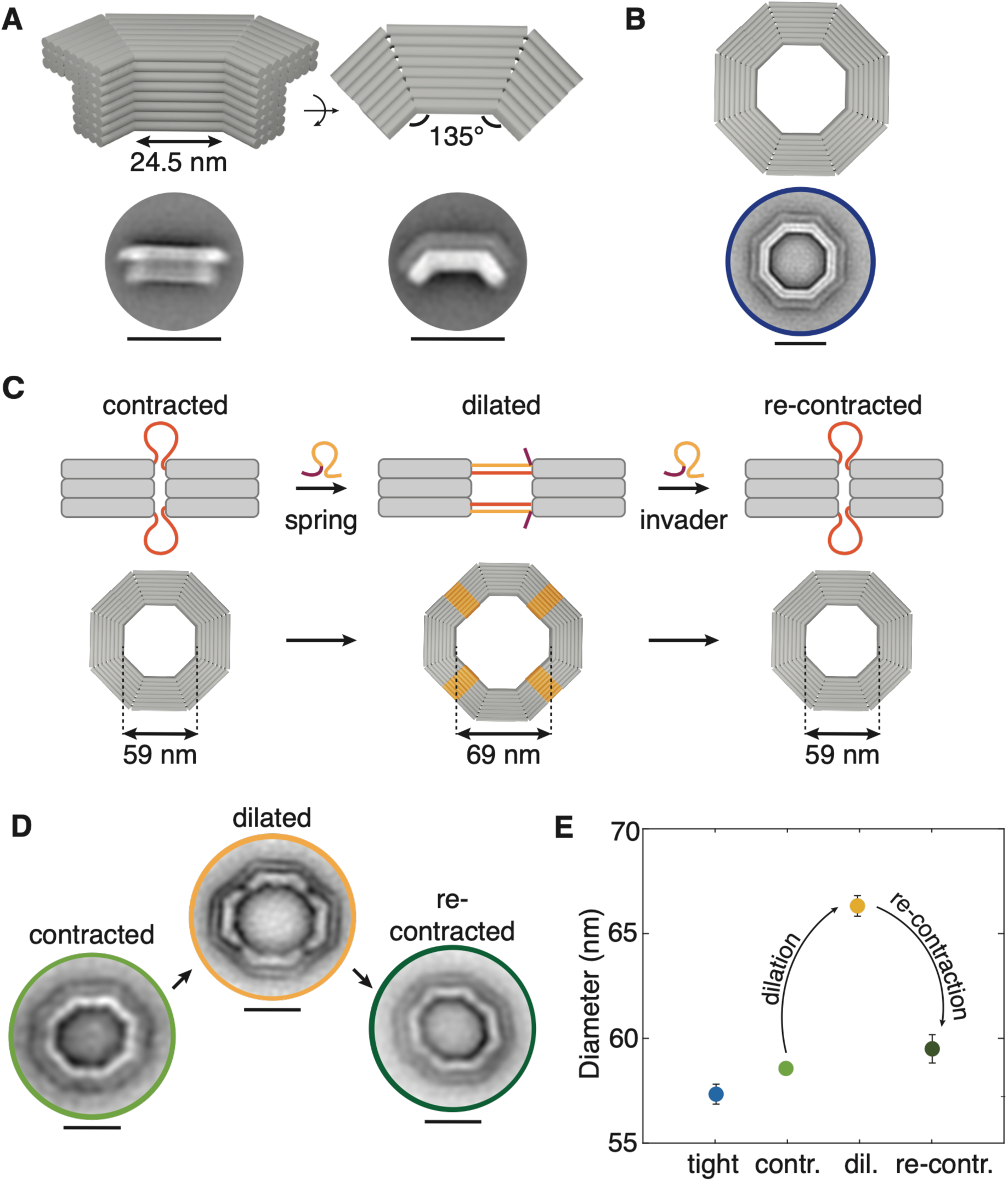
DNA origami nanostructure that size-mimics the NPC. **(A)** DNA origami monomer: Model (top, each cylinder represents a DNA double helix) and 2D TEM class averages (bottom), side and top view. **(B)** Model (top) and 2D class averages (bottom) of the fully formed, tightly bound octagonal nanopore. **(C)** Schematic representation of the dilation/contraction mechanism. Monomers are connected via staples (orange) containing single-stranded regions (left) that can become double-stranded via the addition of complementary spring oligonucleotides (middle), thus expanding the nanopore. These oligonucleotides can be removed by adding complementary invader strands to re-contract the structure to the initial state (right). **(D)** 2D class averages of the dilation/contraction mechanism. Scale bars: 50 nm. **(E)** Diameter of different nanopore states (blue: tightly bound nanopore, green: contracted state, yellow: dilated state, dark green: re-contracted state) as measured by TEM.

As a result, four identical monomers can multimerize in a LEGO-like fashion [44] to form a closed octagon (Fig. 2B) with a designed inner and outer diameter of about 59 nm and 75 nm, respectively. The structure was designed with caDNAno [45] and folded using DNA origami methods [31, 32]. Briefly, a 7560 nt-long scaffold was folded into the desired shape using a set of complementary oligonucleotides. To obtain monomeric structures, the structure was initially prepared by adding oligonucleotides terminating with 3 thymines (poly-Ts) at the interfaces to prevent polymerization. Optimal folding was achieved by annealing the reaction upon slow cooling from 60 °C to 20 °C (1hours/°C) in a buffer containing 18 mM MgCl_2_ (Fig. S3). Transmission Electron Microscopy (TEM) 2D class averages (Fig. 2A, Fig. S4) confirmed that the monomer adopted the desired shape, with an edge length of 23 nm as measured from TEM class averages, closely matching the design. Notably, in the 2D class averages, the flap region is clearly distinguishable from the transmembrane part due to its slightly larger size. Moreover, in the top view (Fig. 2A, right), the flap intensity is lower than the transmembrane region, as it comprises fewer DNA helices.

Four monomers can polymerize to form a complete octagon. To enable the assembly, we folded a variant of the structure without poly-Ts in which the interfaces contained blunt-end helices and scaffold loops (Fig. S2). The blunt-end helices promoted multimerization through **π**-**π** stacking interactions [46, 47], while the remaining helices were bound via connector oligonucleotides, which were complementary to scaffold loops of two neighboring monomers [32]. Of the 44 helices at the interface, 16 were blunt-end helices, while 28 were bound via connector oligos. These connector oligos are fully complementary to the DNA origami scaffold, i.e. we expected the resulting octagons to be composed of tightly bound monomers. The optimal assembly conditions were to incubate the monomers with a 4-fold excess of connector oligos at 40 °C in buffers containing 20 mM MgCl_2_ for at least 5 hours (Fig. S5A).

The fully assembled construct exhibited a clear shift in gels compared to the monomeric structure, while 2D class averages from TEM (Fig. 2B, Fig. S6) showed nanostructures with the desired octagonal shape. Again, the transmembrane part is clearly visible as a region more intense than the surrounding flap. As measured via TEM 2D class averages, the inner and outer diameters of such octagons were 57.3 ± 1.0 nm (mean ± SD; Fig. 2E) and 74.3 ± 0.9 nm, respectively. Surprisingly, the monomers were found to assemble not only into tetramers (octagonal shape) but occasionally (∼10% of cases) also into higher-order structures, such as pentamers (decagonal shape). These structures migrated more slowly than the tetramers in the gels (Fig. S5) and appeared as rings with a larger diameter in TEM micrographs (Fig. S6A). These different conformations indicate a certain degree of flexibility of the monomeric structure, which can adapt to form different structures. However, more than 70% of the structures were correctly assembled octagons, as estimated from particles on TEM images (Fig. S6B), demonstrating the successful design, folding, and assembly of the DNA origami pores.

### Octagon rings can dilate and contract

Designing an octagon that can self-assemble from individual monomers offers not only the advantage of enhancing the structure’s rigidity but also allows to introduce a mechanism to dilate the nanopore, thereby increasing its diameter in a user-defined fashion. This was achieved by connecting the monomers via ‘ss-connector’ oligonucleotides containing a 30-nt stretch that was not complementary to the scaffold (Fig. S7) and which was fully single-stranded without self-complementary sequences to prevent self-interaction. By adding ss-DNA oligonucleotides complementary to these 30-nt stretches, the single-stranded regions were transformed into double-stranded helices, which were designed to push the monomers apart and increase the diameter by approximately 10 nm (Fig. 2C). Hence, the dilated octagons have a designed inner and outer diameters of 69 and 85 nm, respectively.

We were able to self-assemble the monomers into octagons using these ss-connectors. To this end, we prepared the monomers as before and added the ss-connectors during polymerization. Similarly to the tightly bound structure, the optimal assembly conditions were to incubate the monomers with a 4-fold excess of connector oligos at 40 °C in buffers containing 20 mM MgCl_2_ (Fig. S5B). TEM micrographs and 2D class averages showed that the full octagonal construct could be obtained (Fig. 2D, left; Fig. S8), although the inner diameter was slightly (∼1 nm) larger than that of the tightly bound structure, with an average of 58.6 ± 0.2 nm (mean ± SD; Fig. 2E). We attributed this slight increase to the electrostatic and steric repulsion among the single-stranded oligo regions at the interface between the monomers.

Interestingly, we could change the conformation of such octagons to increase their diameters. To achieve larger nanopore sizes, we added DNA oligonucleotides (‘springs’, Fig.2C) complementary to the 30-nt stretches. These constructs showed a slightly lower electrophoretic mobility relative to the contracted nanostructure in gel electrophoresis (Fig. S9), indicating a conformational change. Indeed, TEM micrographs and 2D class averages (Fig. 2D, middle; Fig. S10) revealed clearly dilated octagons due to an increased distance between neighboring monomers, indicating that they were shifted apart. The monomer-monomer distance was visible as a lower-intensity region in the 2D class averages, since 28 of the 44 helices forming the nanopore contained a double-stranded DNA stretch formed by the ss-connectors bound to a spring oligo. The measured inner diameter of the dilated octagon was 66.3 ± 1.0 nm (mean ± SD; Fig. 2E), which is about 8 nm larger than the contracted conformation, while the outer diameter was 86.9 ± 0.8 nm. The 2 nm discrepancy between the designed dilated diameter and the measured one may be due to the structural flexibility of the octagons and interactions with the TEM grid surface.

Importantly, we could also reverse the dilation of the structure, back to a contracted state. As we used spring oligonucleotides that included a 6-nt toehold region that is not complementary to the ss-connectors (Fig. S7B), it was possible to remove them via toehold-mediated strand displacement [48] by adding a fully complementary ‘invader’ strand. Ideally, removing the spring oligos would then revert the octagon to its contracted conformation. To test this for our octagons, we added invader strands to the dilated nanopores. Gel images (Fig. S11) revealed that this construct indeed migrated at the same height as the contracted structure, suggesting that the octagon had reverted to its previous conformation. Moreover, 2D class averages show that the increased distance between the monomers has disappeared (Fig. 2D, right; Fig. S12), with the final structure now closely resembling the contracted one. The inner diameter of the resulting re-contracted structure was 59.5 ± 1.4 nm (mean ± SD), which is very close to the 58.6 nm size of the ss-connected starting structure.

Alternatively, we assembled a nanopore variant substituting the ss-connectors with DNA oligonucleotides that could form hairpins (Fig. S7C). Although the assembly to tetrameric structures had a similar yield as with the ss-connectors (Fig. S5B), most of the tested conditions for the dilation showed lower yields (Fig. S9). This may be due to the fact that spring (and, hence, invader) strands are partially self-complementary, and therefore can form hairpins or hybridize with other copies. Hence, we dropped this approach and decided to use the ss-connectors in all subsequent experiments.

In summary, we built an octagonal nanopore that can reversibly dilate and contract via toehold-mediated strand displacement from a 59 to 66 nm diameter, which represents a 25% change in area.

### Probing the kinetics of octagon dilation and contraction using single-molecule FRET

Next, we studied the dilation and contraction of the octagon at the single-molecule level using Förster Resonance Energy Transfer (smFRET) measurements. smFRET is a powerful method to quantify molecular dynamics based on the distance-dependent energy transfer between fluorescent dyes [49–51]. In solution-based smFRET, fluorescently labelled molecules diffuse in and out of a small observation volume in a confocal microscope, and the signal spikes originating from individual molecules are analyzed. To this end, we added an ATTO-488 (donor)-labelled oligo on one interface and an ATTO-643 (acceptor)-labelled oligo on the other interface for each DNA origami monomer. When monomers assemble into a tetramer, the two dyes on the adjacent monomer interface come into close proximity, enabling FRET upon laser illumination (inset in Fig. 3A). On the other hand, in the dilated octagons the monomer interfaces are expected to be too far apart for efficient energy transfer between the dyes. Notably, each octagon host four FRET pairs.

**Figure 3:**
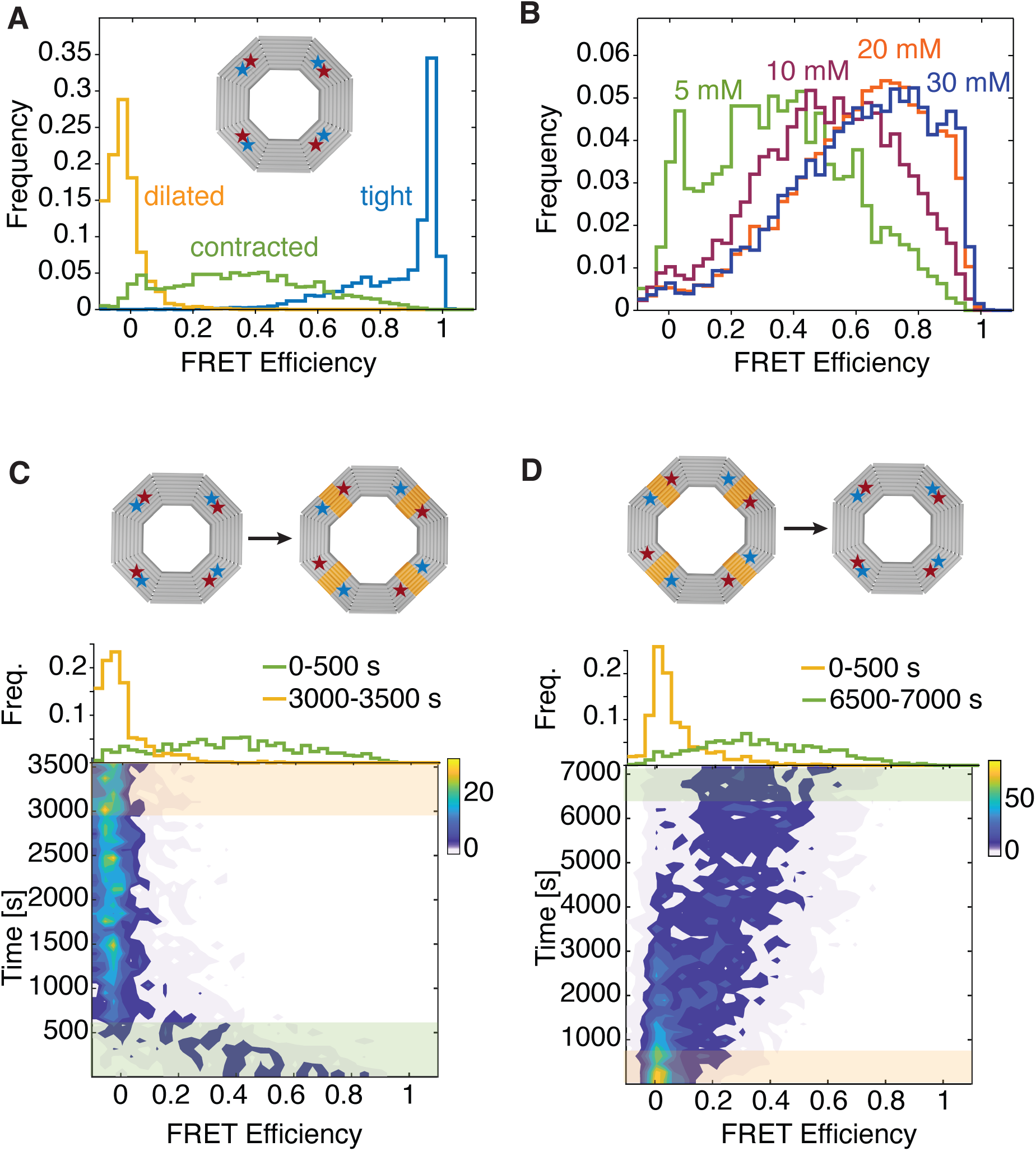
Nanopore dilation and contraction. **(A)** FRET efficiency histograms from single-molecule measurements of the nanopore in different states. Inset: schematic representation of the nanopore, with FRET pairs depicted as stars. **(B)** FRET efficiency histograms from single-molecule measurements of the flexible contracted structure at different MgCl_2_ concentrations, from 5 mM to 30 mM. **(C)** Time course of FRET efficiency when the nanopore changes from a contracted to a dilated state over time. Color from blue to yellow represents the number of events. Green and yellow areas represent the first and the last 500 s, respectively. Green line in the top graph: FRET efficiency of the structure in the first 500 s. Yellow line: FRET efficiency of the structure in the last 500 s. Inset: schematic representation of the nanopore dilation. **(D)** As in (C) but for nanopores changing from a dilated to a contracted state. Yellow line in the top graph: FRET efficiency of the structure in the first 500 s. Green line: FRET efficiency of the structure in the last 500 s.

We first measured the FRET efficiency for the tightly connected octagon. This sample showed a FRET efficiency (*E_FRET_*) close to 1 (Fig. 3A, blue), which is expected due to the close proximity between the dyes in the FRET pair. The FRET efficiency histogram displayed a shoulder down to *E_FRET_* =0.5, which may indicate some structural heterogeneity in the tightly bound octagon, in which the dyes may be further apart than designed.

In contrast, the FRET efficiency of the dilated octagon showed a peak near *E_FRET_* =0 (Fig. 3A, yellow). This signals that the dyes were too far apart for efficient energy transfer, leading to low *E_FRET_* values. This is consistent with the TEM finding that the distance between the monomer interfaces on the dilated octagon is increased by approximately 8 nm, confirming that the nanopore is in a dilated state. Interestingly, the contracted structure bound with ss-connectors showed intermediate FRET efficiencies (broadly distributed around *E_FRET_* ∼0.4; Fig. 3A, green). This indicates that the structure was occupying intermediate states between full contraction and full dilation. As Mg_2+_ ion concentration can affect the **π**-**π** stacking interactions of DNA double helices [44], we conducted smFRET experiments using the ss-connected structure in buffers with Mg^2+^ concentrations ranging from 5 mM to 30 mM. The *E_FRET_* maxima shifted towards higher values with increasing Mg_2+_ concentration (Fig. 3B), but they remained slightly lower than the tightly bound structure. This indicates that the single-stranded connectors may exert a steric effect, creating some distance between the monomers, in alignment with the TEM data, which showed a 2 nm added distance.

Next, we measured the time scale at which the structures change conformation from a contracted to a dilated state. We prepared octagons bound via ss-connectors and added spring oligonucleotides to the solution immediately before smFRET measurements (Fig. 3C). Initially, the curve showing the number of events with a certain FRET efficiency displayed a broad peak at intermediate *E_FRET_* values, as expected for the contracted structure (Fig. 3C, green area and curve). Over time, the structure dilated and the FRET efficiency decreased to near 0. Indeed, comparing the first and last 500 s of the measurements, we obtained spectra (Fig. 3C, yellow and green) similar to those in Fig. 3A. These results demonstrated that the structure switched from a contracted to a dilated state on a timescale of ∼10 minutes. A similar experiment was performed by adding the invader strand to the dilated structure (Fig. 3D) to switch it back to the re-contracted state. The contraction was observed to be much slower than the dilation, with the structures approaching full contraction only after more than 2 hours – consistent with a relatively slow rate of toehold-mediated strand displacement [48].

We thus demonstrated that the octagons can dilate and contract at the single-molecule level, and that conformational changes can be tuned by adjusting ionic strength conditions.

### Octagons integrate into lipid membranes

Next, we showed that it is possible to insert the structure into lipid bilayers, which is nontrivial given their extremely large size (the outer diameter is 74 to 87 nm, as measured for contracted and dilated pores, respectively). For the insertion, we designed cholesterol-modified oligonucleotides with a sequence that is complementary to single-stranded handles protruding beneath the flap (Fig. 4A, left) [43]. Each monomer contained 5 handles; hence, a complete octagon was functionalized with 20 cholesterol-oligos. Positioning the cholesterol tags beneath the flaps resulted in a directional insertion of the nanopore in the membrane. Due to their relatively large size, the octagons were not able to puncture pre-formed lipid bilayers. Hence, we used continuous droplet interface crossing encapsulation (cDICE) [35, 52] to prepare Giant Unilamellar Vesicles (GUVs) containing DNA structures. With this method, DNA origami nanopores were successfully inserted into the membrane during the formation of the lipid bilayer (Fig 4A, right). We validated insertion by confocal microscopy, which revealed colocalization of the fluorescently labelled origami pores with the GUV membrane signal (Fig. 4B). This colocalization was verified across all pore variants used in this study (Fig. S13).

**Figure 4:**
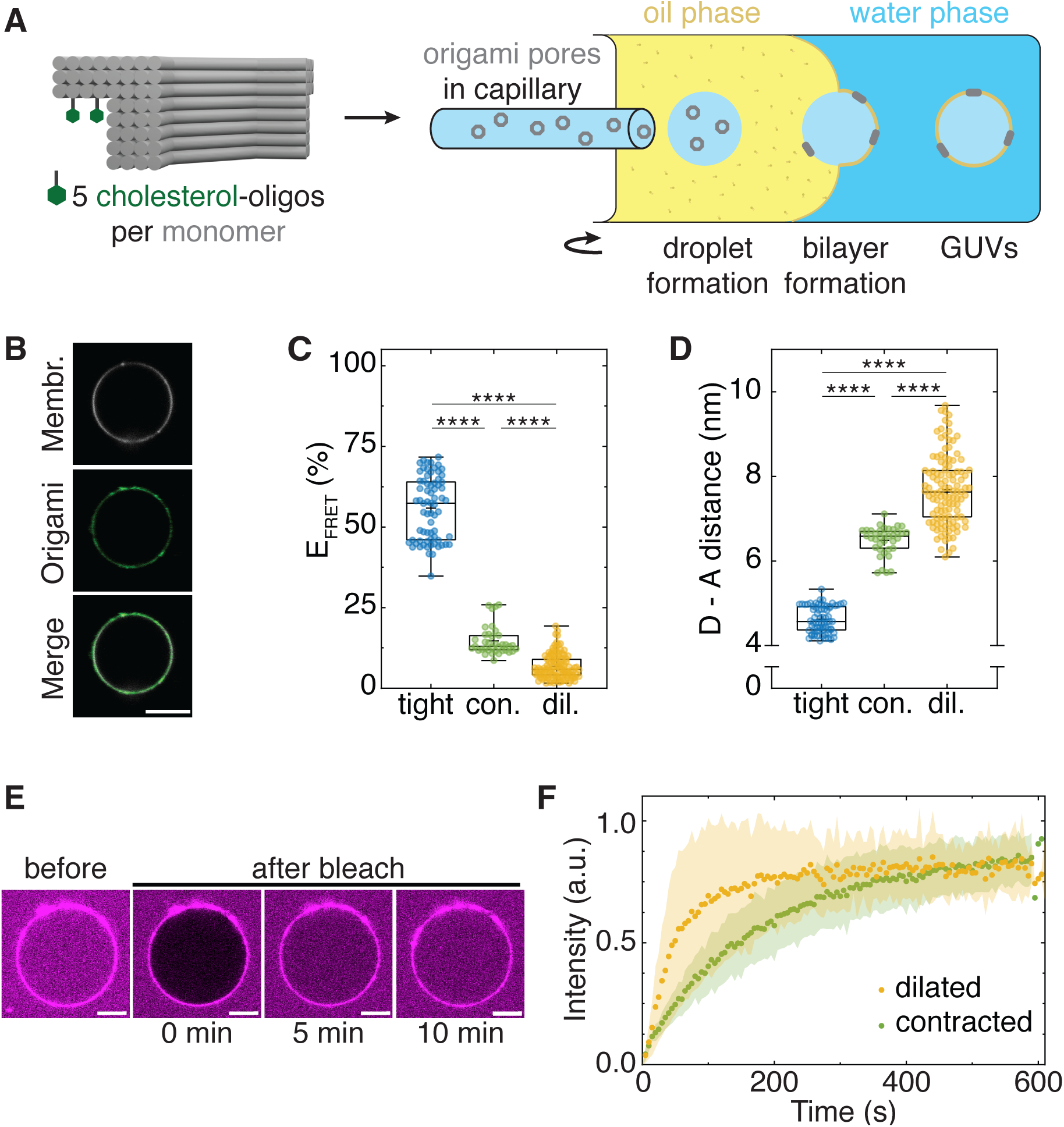
Insertion of origami pores in GUV membranes. **(A)** Schematic representation of the insertion of the origami pores into the membrane of a GUV. Left: monomer with cholesterol anchors indicated in green. Right: GUV preparation via the cDICE method. The origami pores are diluted in the inner solution and droplets are injected into a rotating chamber containing an aqueous and an oil phase. Lipids that are dispersed in the oil phase spontaneously assemble into a monolayer at the water-oil interface. As these droplets move outwards due to the centrifugal force, the lipid-stabilized droplets pass the oil-water interface, resulting in the formation of a second monolayer, and thus a lipid bilayer, around the droplet. **(B)** Representative confocal images showing the insertion of the origami pores into the GUV membrane. Images are acquired at the GUV equatorial plane. Top: ATTO-390 signal (membrane); middle: ATTO-488 signal (origami); bottom: merged channels. Scale bars: 5 µm. **(C)** FRET efficiencies measured for GUVs containing the tight pores (blue; *N* = 4, *n* = 70), pores bound with ss-connectors (green; *N* = 2, *n* = 37), and dilated pores (yellow; *N* = 5, *n* = 118). *N* represents the independent repeats and *n* the individual GUVs. **(D)** Distances between the donor and acceptor of the FRET pair on adjacent monomers. **(E)** Representative confocal images visualizing the mCherry influx in GUVs containing the pores bound with ss-connectors before and after bleaching (directly after, 5 minutes after, and 10 minutes after) (right). Scale bar: 5 µm. **(F)** FRAP curves showing the influx of mCherry in GUVs after photobleaching. Dots represent the average time-dependent intensity I(t), and shaded areas represent the standard deviation across various measurements.

We measured the conformational state of the DNA origami nanopores embedded in the membranes, showing that they maintained their initial configuration. To this end, we performed FRET measurements to determine the inter-monomer spacing in each conformational state. Adjacent monomer interfaces were labelled with ATTO-488 and ATTO-643, and energy transfer was quantified by fluorescence lifetime imaging (FLIM) of the donor dye. A shorter donor lifetime indicated more efficient energy transfer and, consequently, a reduced inter-monomer distance. We first characterized the tightly connected octagon in the lipid bilayers. Consistent with the close proximity of the adjacent monomers in this state, FLIM measurements yielded an intermediate *E_FRET_* of ∼0.6 (Fig. 4C, blue), indicating a spacing of approximately 5 nm between the labelled interfaces (Fig. 4D, blue). For GUVs prepared with the contracted octagon bound with ss-connectors, we observed a lower value of *E_FRET_* (∼0.15; Fig. 4C, green), corresponding to a spacing of approximately 6.5 nm between the interfaces (Fig. 4D, green). GUVs incorporating the dilated octagon yielded an *E_FRET_* close to 0 (Fig. 4C, yellow), indicating that the monomers are interspaced by at least 8 nm (Fig. 4D, yellow), consistent with what we previously observed in the bulk experiments. Taken together, these results demonstrate that the donor fluorescence lifetimes and the corresponding E_FRET_ values obtained for octagons within the lipid bilayer correlate well with the conformational states of the origami structures determined before, confirming that the three distinct pore configurations are preserved upon membrane insertion.

### Octagons allow transmembrane transport of proteins

Next, we demonstrated that the origami structures formed functional open nanopores that facilitate transmembrane transport across the GUV bilayer. Specifically, we studied the transport of the fluorescent protein mCherry across the lipid membranes via fluorescence recovery after photobleaching (FRAP) experiments. To this end, we incubated GUVs containing the different octagon structures overnight with 15 mM mCherry. The fluorescent proteins inside the GUVs were then photobleached, and recovery was monitored over a 10-minute window to quantify the pore-mediated flux across the GUV membrane. For both the dilated and contracted pores, we observed passive diffusion of mCherry into the GUV lumen after photobleaching (Fig. 4E; Fig. S14). This fits expectations, as the hydrodynamic diameter of mCherry (< 5 nm) is much smaller than the inner diameter of all three octagon configurations.

The transport through the larger dilated pores was observed to be faster than through the smaller contracted pores (Fig. 4F), experimentally verifying the existence of larger open pore area. For GUVs containing the contracted octagon bound with ss-connectors, our FRAP measurements yielded a half-time of recovery (t_1/2_) of 106 ± 2 seconds, whereas we obtained a t_1/2_ of 36 ± 1 seconds for GUVs containing the dilated octagon. These results indicate a 3-fold faster recovery for the dilated pores.

These experiments thus demonstrate that the octagon is not only bound to GUVs but can also form open nanopores in lipid bilayers, allowing protein transport through membranes. These octagons, with inner diameters from 58 to 66 nm and outer diameters from 74 to 87 nm, are, to our knowledge, the largest DNA origami nanopores inserted into lipid membranes.

### Building an NPC mimic by Nsp1 functionalization of the octagons

The final step was to functionalize the octagons on their inner lumen with Nups, to thus establish a mimic of the NPC. As a model protein for FG-Nups, we used the well-studied intrinsically disordered region (IDR) of Nsp1 [26]. The Nsp1 IDR contained a single cysteine at the C-terminus to allow for functionalization with an oligonucleotide via cysteine-maleimide chemistry. This oligonucleotide was complementary to a single-stranded anchor protruding from the inner side of the monomers (Fig. 5A). As each monomer contained 30 such handles, a complete octagon can bind up to 120 Nsp1 proteins. The Nsp1 anchor points in the inner surface of the structure have an average distance between each other of ∼6 nm (Fig. S15). Such a configuration allows for a protein density of 240 mg/mL (180 mg/mL for the dilated octagon), which is in line with the estimated human NPC FG-Nup concentrations of 100-300 mg/mL [53–55].

**Figure 5:**
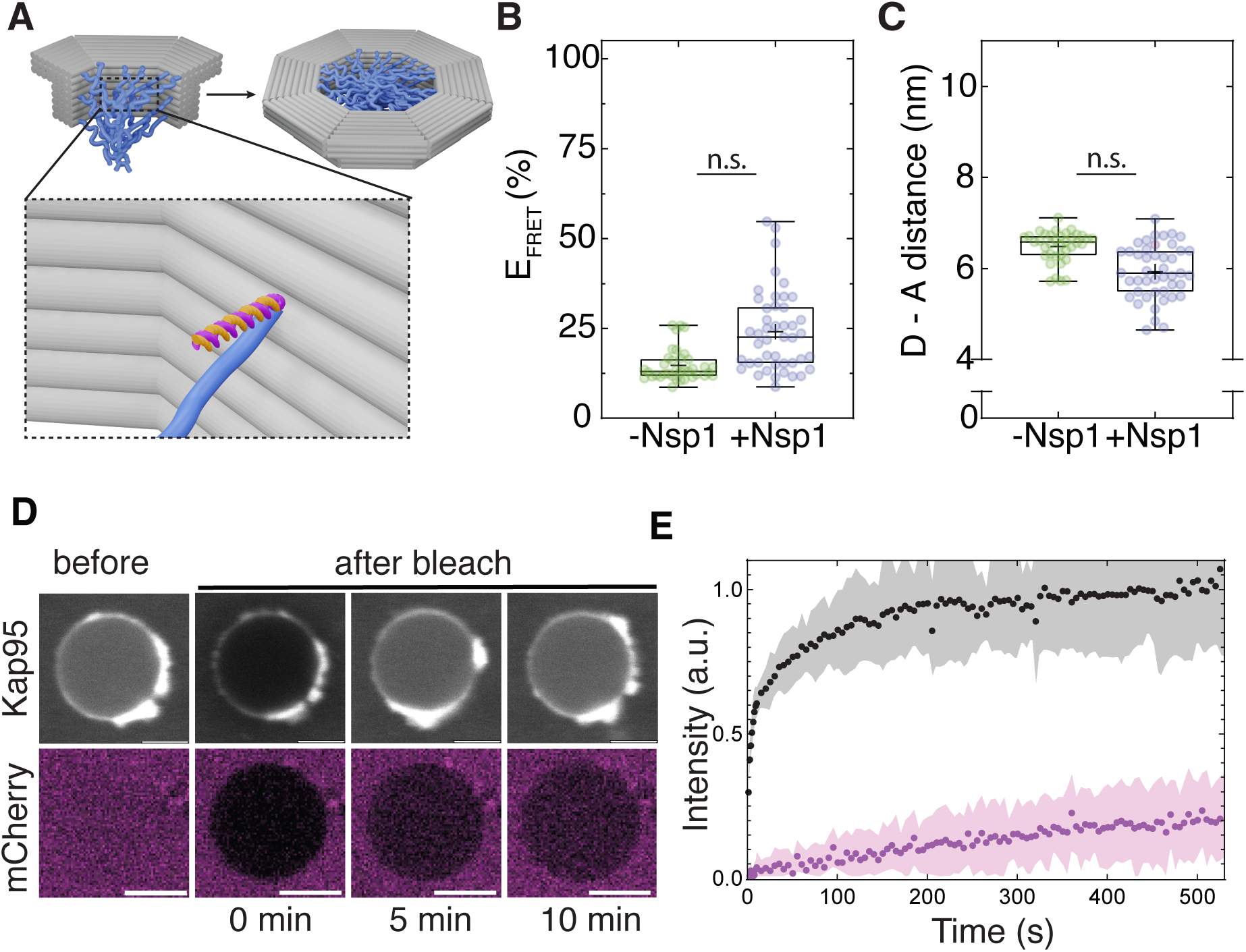
Functionalization of the DNA origami pores with Nsp1 and barrier function. **(A)** Model of the DNA origami octagon (grey) functionalized with Nsp1 proteins (blue). Inset: Nsp1 disordered proteins are functionalized with a single-stranded oligonucleotide (magenta) that is complementary to an anchor oligonucleotide that is protruding from the lumen of the DNA origami nanopore (orange). **(B)** FRET efficiencies measured for GUVs containing the pores bound with ss-connectors with (blue; *N* = 2, *n* = 55) and without (green; *N* = 2, *n* = 37) Nsp1 functionalization. *N* represents the independent repeats and *n* the individual GUVs. **(C)** Distances between the donor and acceptor of the FRET pair on adjacent monomers. The similar data in panels B and C signal that the nanopore size did not significantly change upon Nsp1 functionalization. **(D)** Representative confocal images visualizing Kap95 (top) and mCherry (bottom) are shown before, directly after, 5, and 10 minutes after bleaching. Scale bar: 5 µm. **(E)** FRAP curves showing the influx of Kap95 (black) and mCherry (magenta) in GUVs after photobleaching. Dots represent the average time-dependent intensity I(t) and shaded areas represent the standard deviation.

DNA origami octagons could successfully be functionalized with Nsp1 proteins. After expression and purification, Nsp1 was conjugated with the single-stranded oligonucleotide using cysteine-maleimide chemistry. When examined via gel electrophoresis (Fig. S16), the Nsp1-oligo conjugates showed a clear shift with respect to the protein alone. Adding the Nsp1 to the octagons resulted in a clear electrophoretic shift (Fig. S17A) relative to the bare origami. Such samples were further purified by ultracentrifugation to remove excess oligos and higher-order structures (Fig. S17B).

We thus successfully coated the inner surface of the octagons with Nups. Nsp1 had some effect on the octagon diameter, as we determined by measuring the structure with flexible linkers via smFRET. Interestingly, even in buffers containing 20 mM MgCl_2_, the flexible structure that was designed as the contracted octagon showed lower FRET efficiencies (Fig. S18), similar to the dilated structure in Fig. 3A. We speculate that Nsp1 in these specific buffer conditions act as a bulky polymer brush in the nanopore lumen, pushing the monomers apart.

We next investigated the effect of Nsp1 attachment on the conformational state of the octagon upon membrane insertion. GUVs with Nsp1-conjugated octagon assemblies were prepared using the cDICE method. We validated insertion of the conjugated structure into the GUV membrane via confocal microscopy as we observed the colocalization of the membrane signal and that of the fluorescently labelled origami structure (Fig. S13). To determine the inter-monomer spacing, we performed FLIM measurements following the same approach described earlier. The Nsp1-conjugated contracted octagon yielded an intermediate *E_FRET_* value of ∼0.25 (Fig. 5B), corresponding to an inter-monomer spacing of approximately 6 nm (Fig. 5C). This is close to the value of 6.5 nm we found for the unconjugated contracted octagon bound with ss-connectors in the GUV membranes.

We validated that these octagon structures conjugated with Nsp1 proteins formed functional semi-permeable nanopores in the GUV membranes. To this end, we studied the mCherry diffusion into the GUV lumens via the contracted Nsp1-coated octagons as monitored by FRAP measurements described earlier (Fig. 5D). We observed that mCherry could passively diffuse into the GUV lumen as the vesicles were filled after overnight incubation with the fluorescent proteins. Notably, for GUVs containing the conjugated octagon structure with Nsp1, we observed a t_1/2_ of 360 ± 42 seconds (Fig. 5E), which is ∼3.5 times slower compared to the contracted octagon without Nsp1, and about 10 times slower compared to the dilated pore without Nsp1. This observation indicates that Nsp1 forms a permeability barrier that is restricting the flux of mCherry through the pores. mCherry is a relatively small protein with its 27 kDa mass, and accordingly it may penetrate the Nup barrier via passive diffusion [8, 9]. To test whether bigger macromolecules are prevented from crossing the lipid membrane, we inserted DNA origami octagons coated with Nsp1 in GUVs, and added FITC-dextran in the outer solution (Fig. S19). Notably, we used 150 kDa dextran, which corresponds to a diameter of gyration of 20 nm [35, 56], smaller than the nanopore diameter. Even after overnight incubation at room temperature, the interior of the vesicles barely showed any fluorescent signal, indicating that these big FITC-dextran macromolecules are not able to cross the Nsp1 barrier.

Finally, to investigate whether selective transport can be mediated by the Nsp1-functionalized nanopores, we monitored the translocation of Alexa647-labelled Kap95 into GUV lumens by FRAP measurements as described earlier (Fig. 5D-E). Kap95 entered the GUV lumens rapidly with a t_1/2_ of 30 ± 1 seconds. Notably, this is 12 times faster than mCherry, which passively diffuses through the pores with a t_1/2_ of 360 seconds. This pronounced difference in translocation rate between the transport receptor Kap95 and the inert small protein mCherry demonstrates that the Nsp1 meshwork confers selective permeability as it effectively restricts passive diffusion while actively facilitating the passage of Kap95.

## Discussion and conclusions

In this work, we designed, folded, assembled, and tested a tetrameric octagonal origami nanopore whose inner diameter can be tuned from ∼57 to ∼66 nm sizes, comparable to the NPC dimensions in contracted and dilated states. We showed that these octagons can reversibly be dilated and contracted. TEM measurements showed a change in the nanostructure’s inner diameter of 8 nm, corresponding to an area change of 25%. We showed that the dilation from a contracted state is relatively fast (happening on a timescale of ∼10 minutes) but re-contraction was slower (timescale of hours), likely because of the slow kinetics of toehold-mediated strand displacement [48]. In the future, alternative actuation mechanisms may be implemented to achieve a faster dilation and contraction. For instance, photoswitchable moieties such as _cnv_K [57] or azobenzene [58] could be incorporated at the monomer interfaces, offering a route to optically regulate conformational changes. We could insert both the contracted and the dilated DNA origami octagon into GUVs, yielding the largest DNA origami nanopore inserted into lipid membranes to date. Both conformational states were stable after insertion, with dilated nanopores exhibiting a much larger diameter than contracted ones, in good agreement with the measurements in solution.

To mimic the selective transport properties of the NPC, we functionalized the nanopores with the intrinsically disordered domain of Nsp1 and inserted these constructs into GUVs. Intriguingly, the effect of Nsp1 was different in solution and in GUV experiments. In solution, the presence of Nsp1 led to an increase in the distance between the monomers. In contrast, when the same Nsp1-functionalised nanopores were embedded in GUVs, the distance between the monomers decreased. This suggests that membrane anchoring restricts the nanopore dilation, although differences in buffer composition between the experiments with free DNA origami nanopores and in GUV could also contribute. Interestingly, the presence of Nsp1 significantly restricted the flux of the fluorescent reporter mCherry through the pores, indicating that it formed a permeability barrier. In contrast, the transport receptor Kap95 strikingly translocated 12 times faster through the Nsp1-functionalized nanopores, which is a direct functional readout of Nsp1-mediated selectivity, reconstituted here in a minimal bottom-up system.

Overall, we have established a biomimetic DNA origami nanopore inspired by the NPC architecture, with a tunable inner diameter and regulated NTR transport selectivity properties. Notably, this is the first time that a DNA origami nanostructure coated with intrinsically disordered Nups and embedded in lipid bilayers has been used to study NTR transport in GUVs. In fact, previous research on NPC DNA origami biomimetic systems either lacked embedding of the nanostructures in membranes [36, 39, 40] or used dextran instead of biologically relevant NTRs as cargo [38]. This system thus takes us a step closer to constructing an artificial nucleus from the bottom up. The modular design and the directional insertion into membranes, which is achieved by positioning cholesterol tags under the horizontal flap region of the octagons, open the possibility to build more complex biomimetic systems. In particular, the octagon could be functionalized with multiple different types of nucleoporins that are placed in precise and pre-defined positions to create specific gradients in FG-Nups composition. Such systems will constitute unique platforms to dissect the NPC transport properties from a bottom-up perspective and allow investigating the relationships between several Nup types in relation to the nanostructure diameter.

Beyond studying its biological counterpart, our NPC biomimetic system offers new opportunities for the field of synthetic biology. For instance, it could be used as a synthetic channel to selectively regulate the transport of biomolecules in artificial cells, thereby contributing to developing controllable communication and transport interfaces in bottom-up systems.

## Methods

### Design of scaffolded DNA origami nanostructures

DNA origami nanostructures were designed with caDNAno [45]. The scaffold was 7560 nucleotides long.

### DNA origami folding conditions

The folding reaction for the monomeric structures contained p7560 scaffold (tilibit GmbH) at a final concentration of 40 nM, oligonucleotides (IDT Integrated DNA Technologies) at 200 nM each, and folding buffer containing 1 mM EDTA, 5 mM NaCl, 10 mM Tris-HCl (pH 7.5), and 18 mM MgCl_2_. The folding reaction was heated at 65 °C for 15 min, followed by a temperature ramp from 60 °C to 20 °C (1 °C per hour) in a PCR machine (Biorad). The reaction was then left in the fridge before other sample preparation steps.

### DNA origami purification via ultrafiltration

Excess oligonucleotides were removed via ultrafiltration [32, 59] using Amicon Ultra 0.5 mL Ultracel filters 50K, using 1xFoB buffer (10 mM Tris-HCl, 1 mM EDTA, 5 mM NaCl) with 5 mM MgCl_2_. All the centrifugation steps were performed at RT, at 10k g for 3 min. First, 500 µL of buffer was centrifuged through the filter. After discarding the permeate, 50 to 100 µL of sample was added to the filter. The solution was topped up with buffer to 500 µL and centrifuged again. After discarding the permeate again, 450 µL of buffer was added and centrifuged. This step was repeated 5 times in total before retrieving the sample. To retrieve the purified sample, the filter insert was placed upside down in a new tube and centrifuged for 5 min. Ultrafiltration was performed after folding, assembly, and functionalization to remove all the excess oligonucleotides.

### DNA origami purification via ultracentrifugation

A series of buffers were prepared using 1x TE supplemented with 5 mM MgCl₂ and varying concentrations of glycerol (60% to 30%, in 5% steps). In glass tubes, 200 µL of the 60% glycerol buffer was carefully pipetted to create the bottom layer, followed by 400 µL of the other buffer solution in decreasing glycerol concentrations. The gradients were then incubated overnight at 4°C. Approximately 200 µL of the sample was added on top of the established gradients. If there was remaining space, a 15% glycerol buffer was added. The ultracentrifuge (Beckman Coulter Optima_TM_ L-90K)) run at 4°C for 45 minutes at 40,000 rpm. To retrieve samples, aliquots of 200 µL were pipetted from the top of the tube, ensuring that the pipette tip did not penetrate below the surface of the solution. Agarose gel analysis was performed to identify the sample aliquots.

### DNA origami assembly conditions

For assembly, polymerization oligonucleotides were added in 4x excess with respect to the purified monomer and the MgCl_2_ concentration was adjusted to 20 mM. The reaction was incubated overnight at 40 °C and then purified via ultrafiltration.

### DNA origami functionalization with cholesterol oligonucleotides

Cholesterol oligonucleotides (IDT Integrated DNA Technologies) were heated to 50 °C for 15 min before use and added at 5x excess to each handle (5 handles/monomer) in 1xFoB with 5 mM MgCl_2_. The solution was incubated at RT for at least 4h and then purified via ultrafiltration.

### Nsp1 expression and purification

The Nsp1 sequence used in this study is: MHHHHHHHHHHGSGENLYFQGTSMGNFNTPQQNKTPFSFGTANNNSNTTNQNSSTGAGAFGTGQSTFGFNNSAPNNTNNANSSITPAFGSNNTGNTAFGNSNPTSNVFGSNNSTTNTFGSNSAGTSLFGSSSAQQTKSNGTAGGNTFGSSSLFNNSTNSNTTKPAFGGLNFGGGNNTTPSSTGNANTSNNLFGATANANKPAFSFGATTNDDKKTEPDKPAFSFNSSVGNKTDAQAPTTGFSFGSQLGGNKTVNEAAKPSLSFGSGSAGANPAGASQPEPTTNEPAKPALSFGTATSDNKTTNTTPSFSFGAKSDENKAGATSKPAFSFGAKPEEKKDDNSSKPAFSFGAKSNEDKQDGTAKPAFSFGAKPAEKNNNETSKPAFSFGAKSDEKKDGDASKPAFSFGAKPDENKASATSKPAFSFGAKPEEKKDDNSSKPAFSFGAKSNEDKQDGTAKPAFSFGAKPAEKNNNETSKPAFSFGAKSDEKKDGDASKPAFSFGAKSDEKKDSDSSKPAFSFGTKSNEKKDSGSSKPAFSFGAKPDEKKNDEVSKPAFSFGAKANEKKESDESKSAFSFGSKPTGKEEGDGAKAAISFGAKPEEQKSSDTSKPAFTFGAQKDNEKKTEC Nsp1 was kindly provided by the Görlich lab (Göttingen, Germany) in lyophilized form and was resuspended in denaturing buffer. The plasmid was first described in [53].

### KapG5 expression and purification

The Kap95 sequence used in this study is: GPASVGSMSTAEFAQLLENSILSPDQNIRLTSETQLKKLSNDNFLQFAGLSSQVLIDENTKLEGRILAALTLKNELVSKDSVKTQQFAQRWITQVSPEAKNQIKTNALTALVSIEPRIANAAAQLIAAIADIELPHGAWPELMKIMVDNTGAEQPENVKRASLLALGYMCESADPQSQALVSSSNNILIAIVQGAQSTETSKAVRLAALNALADSLIFIKNNMEREGERNYLMQVVCEATQAEDIEVQAAAFGCLCKIMSLYYTFMKPYMEQALYALTIATMKSPNDKVASMTVEFWSTICEEEIDIAYELAQFPQSPLQSYNFALSSIKDVVPNLLNLLTRQNEDPEDDDWNVSMSAGACLQLFAQNCGNHILEPVLEFVEQNITADNWRNREAAVMAFGSIMDGPDKVQRTYYVHQALPSILNLMNDQSLQVKETTAWCIGRIADSVAESIDPQQHLPGVVQACLIGLQDHPKVATNCSWTIINLVEQLAEATPSPIYNFYPALVDGLIGAANRIDNEFNARASAFSALTTMVEYATDTVAETSASISTFVMDKLGQTMSVDENQLTLEDAQSLQELQSNILTVLAAVIRKSPSSVEPVADMLMGLFFRLLEKKDSAFIEDDVFYAISALAASLGKGFEKYLETFSPYLLKALNQVDSPVSITAVGFIADISNSLEEDFRRYSDAMMNVLAQMISNPNARRELKPAVLSVFGDIASNIGADFIPYLNDIMALCVAAQNTKPENGTLEALDYQIKVLEAVLDAYVGIVAGLHDKPEALFPYVGTIFQFIAQVAEDPQLYSEDATSRAAVGLIGDIAAMFPDGSIKQFYGQDWVIDYIKRTRSGQLFSQATKDTARWAREQQKRQLSLLPETGGG Kap95 was purified as described previously [54] and C-terminally labeled with AZDye647, which is structurally identical to AlexaFluor647, using sortase-mediated ligation [60] as described previously [26].

### DNA origami functionalization with Nsp1-oligonucleotides

Lyophilized amine-functionalized oligonucleotides were resuspended in ddH₂O to a stock concentration of 1 mM. To ensure complete dissolution, oligonucleotides were first incubated at 30°C with shaking at 400 rpm for 30 minutes, followed by a second incubation at 45°C and 700 rpm for 30 minutes. Oligonucleotide concentration was subsequently verified by UV absorbance measurement using a NanoDrop spectrophotometer. The heterobifunctional crosslinker Sulfo-SMCC (sulfosuccinimidyl-4-(N-maleimidomethyl)cyclohexane-1-carboxylate, ThermoFisher Scientific) was equilibrated to room temperature prior to use and dissolved in ddH₂O to a final concentration of 9 mM. The conjugation reaction was performed in 100 mM HEPES buffer (pH 7.5) at room temperature for 2 hours, using a 10-fold molar excess of Sulfo-SMCC relative to the amine-oligonucleotide. Following the reaction, unreacted NHS-ester groups were quenched by addition of Tris buffer at a 10-fold molar excess over Sulfo-SMCC. The maleimide-functionalized oligonucleotide was immediately purified by ethanol precipitation to remove excess crosslinker and reaction byproducts. Briefly, 3 M sodium acetate was added in 1/10 of total reaction volume, followed by addition of two volumes of ice-cold ethanol (100%, reaction grade, pre-cooled at −20°C for at least 30 minutes). The mixture was incubated at −20°C for 30 minutes to allow precipitation, then centrifuged at maximum speed at 4°C for 45 minutes. The supernatant was carefully removed with a pipette, and the pellet was washed once with ice-cold 70% ethanol (volume equivalent to the original reaction volume). After removal of the wash supernatant, the pellet was dried in a desiccator at 30°C for approximately 20 minutes or until complete ethanol evaporation. For conjugation to the protein of interest, the dried oligonucleotide pellet was directly resuspended in the protein solution, assuming complete oligonucleotide recovery in the pellet for stoichiometric calculations. The reaction was carried out at room temperature for 1 hour at 400 rpm, followed by overnight incubation at 4°C. The reaction was purified via gel filtration with a S200 column in a buffer containing 6M GuHCl. Purified Nsp1-oligonucleotide conjugates were added to 1.5x excess to origami binding sites (30 binding sites per monomer = 120 binding sites per octagon) overnight at room temperature. The buffer was adjusted to contain 2 M GuHCl and 20 mM MgCl_2_.

### DNA origami gel electrophoresis

Samples were electrophoresed in gels containing 2% agarose, 0.5xTBE (Tris-borate-EDTA), and 5 mM MgCl_2_ for 1.5 (monomers) to 2 hours (tetramers) at 90 V bias voltage in a gel box. Gels that were running for more than 1.5 hours were cooled in an ice bath. The gels were imaged in a gel imager (BioRad Universal Hood II Gel Doc Imaging System). Image processing was performed with FIJI [61].

### Protein gel electrophoresis

Samples were electrophoresed in a pre-cast 4-12% BioRad gel in 1xMES buffer for 30 min at 200 V in a vertical gel box. After the run, the gels were rinsed in water, stained with Instant Blue Coomassie (Sigma Aldrich) for 30 min, and de-stained overnight in water. The gels were imaged in a gel imager (BioRad Universal Hood II Gel Doc Imaging System). Image processing was performed with FIJI [61].

### TEM

A carbon Formvar grid (Electron Microscopy Sciences) was glow-discharged (1 min, 45 mA) before use. The sample (5 µL, 5-20 mM MgCl_2_) was pipetted onto the grid. The incubation time depended on the sample concentration (30 s for ∼50 nM, up to 10 min for ∼1 nM concentrations). The excess sample was blotted with a filter paper. A 5 µL droplet of stain solution (2% uranyl acetate) was pipetted onto the grid and immediately blotted. Another 5 µL droplet of stain was pipetted onto the grid, incubated for 30 s, and blotted away. The grid was left to dry in air for minimum 10 min before imaging on a Jeol FSC3200 microscope. Class averages were computed using Relion 3.0 [62].

### Single-molecule FRET experiments in solution

Samples were diluted to a final concentration of 300 pM in 1xFoB buffer containing 1 mM Trolox and 20 mM MgCl_2_ unless otherwise specified. Samples were pipetted into an µ-Slide 15-well 3D (ibidi) and measured using a MicroTime 200 microscope (PicoQuant GmbH, Germany) using the SymPhoTime software. The measurements were processed using PAM analysis software [63] in MATLAB.

### GUV formation

GUVs were formed using cDICE [52]. The inner buffer contained 50 mM TRIS, 1 mM EDTA, 5 mM NaCl, 20 mM MgCl_2_, and 15 µM Dxt (MW=2 MDa). This buffer was used to dilute the DNA origami sample to 200 µL at a final concentration between 4-10 nM. The outer buffer was obtained by adding glucose into ddH_2_O until the osmolarity was the same as the inner buffer. The osmolarities of the inner and outer buffers were measured using a freezing-point Osmometer (Gonotec Osmomat 3000; Bruker). For influx experiments, proteins were diluted in a buffer containing 25 mM TRIS, 20 mM MgCl_2_, and NaCl to have the same osmolarity as the other buffers. The lipid mixture contained 99.5 mol% 1,2-dioleoyl-sn-glycero-3-phosphocholine (DOPC; Avanti Polar Lipids) and 0.5 mol% 1,2-dioleoyl-sn-glycero-3-phosphoethanolamine (DOPE) labelled with ATTO390 (AttoTec) at a nominal concentration of 0.5 mg/mL. The lipid mixture was prepared in a glass beaker with chloroform-cleaned glass syringes and dried using N_2_. In a glovebox with a dehumidified N_2_ environment, 1.3 mL mineral oil was mixed with 5.7 mL silicon oil. Lipids were resuspended in 50 µL chloroform and 400 µL decane. The oil mixture was added drop-by-drop to the lipid mixture while vortexing. Afterwards, the whole mixture was vortexed for 2 min and sonicated for 15 min in an ice bath. For cDICE, the syringe and capillary were first cleaned 1x with 1 mL ddH_2_O and 2x with 1 mL inner buffer. 500 µL outer buffer was added to the rotating chamber, followed by 5.5 mL of the lipid-in-oil mixture. The origami sample was diluted in the inner buffer and collected into the syringe. The capillary was inserted 2 mm into the lipid-in-oil mixture, and the syringe pump was started with a flow of 20 µL/min. When the capillary was empty, it was immediately removed from the rotating chamber, which was left spinning for another 30 s. The chamber was then positioned vertically on a surface to accumulate the GUVs at the bottom. 200 µL of sample was collected for imaging and pipetted into µ-Slides 8-Well high bioinert (ibidi).

### Confocal microscopy

Imaging was performed on an inverted Leica Stellaris 8 FALCON laser scanning confocal microscope using LasX software (version 4.8.2). Samples were illuminated using a 63x glycerol immersion objective (HC PL APO CS2 63X/1.30, Leica). Atto390-DOPE, used to label the GUV membrane, was excited at 405 nm using a solid-state diode laser. Molecular brightness was optimized by adjusting the correction ring of the objective prior to the measurement. The emission light was passed through a 103 µm pinhole and collected between 415 – 480 nm by a HyD X detector operated in digital mode with an internal gain of 25. Confocal images of 512 × 512 pixels with a pixel dwell time of 2.8 µs were acquired at the GUV midplane at a scan speed of 400 Hz.

FLIM-FRET measurements were performed using the same Stellaris 8 FALCON confocal microscope. Atto488 was excited at a wavelength of 497 nm using a white light laser at a repetition rate of 20 MHz. Brightness was optimized by adjusting the objective correction collar prior to each experiment. To ensure the acquisition of enough photons per pixel to obtain reliable decay fits without affecting the photon rate per pulse, the number of line repetitions per image was adjusted. The emission light was passed through a 103 µm pinhole and collected between 507 - 640 nm using a HyD X detector operated in digital mode with an internal gain of 100.

FRAP measurements were performed using the same confocal microscope as described above, operated using the FRAP wizard in the LasX software. A series of confocal images consisting of 5 pre-bleach frames, 2 bleach frames, and 100 post-bleach frames was acquired at the equatorial plane of GUVs with a 10 s interval. To visualize mCherry, the pre/post bleach frames were obtained at an excitation wavelength of 587 nm, and the emission light was passed through a 103 µm pinhole and collected between 597 – 700 nm using a HyD S detector operated in analog mode with an internal gain of 30. To visualize Alexa647-labelled Kap95, the pre/post bleach frames were obtained at an excitation wavelength of 646 nm, and the emission light was passed through a 103 µm pinhole and collected between 656 – 750 nm using a HyD S detector operated in analog mode with an internal gain of 50. A circular region with a radius of 3 μm in the GUV lumen was bleached using the *fly mode*, *zoom-in*, and *background to zero* settings in the FRAP wizard.

### Fluorescence lifetime analysis

Fluorescence lifetime analysis was performed using the single molecule wizard of the Leica LasX software. The Atto 488 signal at the equatorial plane of each GUV was manually selected. Photon arrival times were fitted with an n-exponential reconvolution function with n = 1 for samples containing origami pores labelled with only the donor, and n = 2 for all other samples. The instrument response function (IRF) generated by the software was used. The amplitude-weighted lifetimes were used for calculating FRET efficiencies. Donor (Atto488) decay functions are approximated by a bi-exponential model:

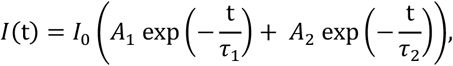

with I(t)the intensity at time t, I_0_ the intensity at time t = 0, and A_1_ and A_2_ pre-exponential factors associated with *τ*_1_ and *τ*_2_. By using this model, amplitude-weighted lifetimes are given by:

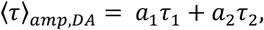

where a_1_ and a_2_ are the amplitudes of the lifetime components *τ*_1_ and *τ*_2_ and a_1_ + a_2_ = 1. The resulting FRET efficiencies were then calculated as follows:

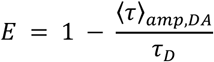

To convert the FRET efficiencies into donor - acceptor distances we used

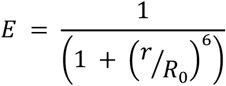

With r the donor – acceptor distance and R_0_ the Förster distance (4.8 nm for the FRET pair used here).

### FRAP analysis

To determine the fluorescence recovery times for mCherry and Atto647N-Kap95 after photobleaching, we analyzed the time-dependent intensity I(t) during recovery in the bleached region throughout fluorescence recovery [64]. We corrected for photofading during the acquisition of post-blech series by also measuring the fluorescence over time in a reference region away from the bleaching region. Then, we scaled the fluorescence intensity into a 0–1 range and we computed the normalized intensity I_n_(t) by dividing the scaled I(t) by the prebleach intensity *Ii*.

### Statistics

Values for E_FRET_, the D – A distances, and the recovery curves were plotted in Graphpad Prism (version 10.6.1). All statistical analysis were performed in the same software. Prior to statistical comparison, the data were evaluated for normality and homogeneity of variance (Shapiro-Wilk test). As these conditions were not met, we used a non-parametric Kruskal–Wallis ANOVA test followed by a Dunn’s multiple comparisons test to evaluate the statistical significance between the samples.

## Data availability

The raw data and processed data for reproducing the figures are deposited in Zenodo and will be made publicly available upon publication of the manuscript. The DNA nanostructure design will be uploaded to nanobase.org upon publication of the manuscript.

## Acknowledgements

We thank Yuchen Yan and Owen Luflos for supporting experiments; Eli van der Sluis and Ashmiani van der Graaf for support with Nsp1 protein; Nils Klughammer and Jaco van der Torre for insightful discussions. We acknowledge funding by the NWO-XL grant OCENW.GROOT.2019.068 and by the NWO-XS grant OCENW.XS23.3.049 of the Dutch Research Council (NWO).

## Author contributions

Conceptualization: C.D., E.B.; Investigation and Formal analysis: E.B., B.v.H, A.B., S.W.; smFRET software: A.B.; Methodology: E.B., B.v.H., A.B.; Writing - Original Draft: E.B., B.v.H. Writing - Review & Editing: E.B., B.v.H, C.D., A.B., S.W.; Funding acquisition: C.D., E.B.; Supervision: C.D.

